# Eukarion-134 attenuates endoplasmic reticulum stress-induced mitochondrial dysfunction in human skeletal muscle cells

**DOI:** 10.1101/2020.06.10.143958

**Authors:** Anastasia Thoma, Max Lyon, Nasser Al-Shanti, Gareth A Nye, Robert G Cooper, Adam P Lightfoot

## Abstract

Maladaptive endoplasmic reticulum (ER) stress is associated with modified reactive oxygen species (ROS) generation, altered mitochondrial bioenergetics, and oxidative damage; and is postulated as a potential mechanism involved in the underlying muscle weakness experienced by patients with myositis, an acquired autoimmune neuromuscular disease. In this study, we investigate the impact of ROS generation in an *in vitro* model of ER stress in skeletal muscle, using the ER stress inducer tunicamycin (24 hours) in presence or absence of a superoxide dismutase/catalase mimetic Eukarion (EUK)-134. ER stress activation, ROS generation, mitochondrial function, biogenesis, morphology and dynamics (fusion/fission) were examined. Tunicamycin induced maladaptive ER stress, validated by stimulation of GRP94, GRP78, CHOP, XBP-1, ERDJ4, and GADD34, which were mostly mitigated by EUK-134 at transcriptional level. ER stress triggered mitochondrial unfolded protein response and promoted mitochondrial dysfunction, described by substantial loss of mitochondrial membrane potential, as well as reduction of respiratory control ratio, reserve capacity, phosphorylating respiration, and coupling efficiency, which was ameliorated by EUK-134. ROS-mediated biogenesis and fusion of mitochondria was evident in presence of tunicamycin, which however, had high propensity of fragmentation, accompanied by upregulated mRNA levels of fission-related markers. Increased cellular ROS generation oxidative stress was observed in response to ER stress that was ameliorated in the presence of EUK-134, even though no changes in mitochondrial superoxide were noticeable. These findings suggest that targeting ROS generation using the superoxide dismutase/catalase mimetic EUK-134 can amend aspects of ER stress-induced changes in mitochondrial dynamics and function. Overall, this study suggests that in instances of chronic ER stress, such as in myositis, quenching ROS generation may be a promising therapy for muscle weakness and dysfunction.

## INTRODUCTION

The endoplasmic reticulum (ER) is a specialised organelle, which is the key site of protein folding in the cell. The reducing environment of the ER contributes to the high fidelity needed to correctly fold newly synthesised peptides and proteins into their biological active conformation [1, 2]. The accumulation of misfolded or aggregated proteins within the ER, termed ER stress, initiates the unfolded protein response (UPR), a ubiquitously expressed network of cellular processes, responsible for restoring protein homeostasis [3]. The UPR induces ER-associated degradation via the ubiquitin proteasome pathway (26S proteasome) to facilitate clearance of misfolded proteins, inhibits protein assembly via translation attenuation, and increases protein folding capacity via chaperones release (glucose-regulated protein (GRP) 78 and 94). However, failure of establishing protein homeostasis and sustained ER stress lead to UPR-induced cell death, through caspase 12 (autophagy) and cholesterol oxidase-peroxidase C/EBP homologous protein (CHOP) activation (apoptosis) [4, 5, 6].

It is well established that chronic or prolonged activation of the ER stress response is implicated in a wide range of diseases, such as neurodegenerative diseases (e.g., Alzheimer’s disease), chronic metabolic diseases (e.g., diabetes), and muscle diseases (e.g., Duchenne muscular dystrophy and myositis) [7, 8, 9, 10]. The observation that aberrant activation of the ER stress pathway in the muscles of patients with myositis, a rare neuromuscular disorder of autoimmune origin, has directed our research interest at the understanding of the downstream mechanisms of ER stress in skeletal muscle.

The close approximation of ER and mitochondria permits bi-directional crosstalk via the mitochondrial-associated ER membranes and Ca^2+^ signalling, highlights the potential impact of the sarcoplasmic reticulum (ER in skeletal muscle) on mitochondrial function. Reactive oxygen species (ROS) are generated as an upstream and a downstream component of the UPR pathway. Hydrogen peroxide (H_2_O_2_) is generated via the thiol/disulphide (-SH/-SS) exchange mechanism during protein folding in the ER, as well as from superoxide (O_2_^·-^) generated from mitochondrial complexes I and III following Ca^2+^ influx. In addition to H_2_O_2_, peroxynitrite (ONOO^-^) is also generated as a by-product of the reaction between O_2_^-^ and nitric oxide (NO), which its generation depends on Ca^2+^-stimulated nitric oxide synthase [6, 11]. Prolonged activation of the ER stress response can influence mitochondrial function, resulting in aberrant ROS generation, from both the ER lumen and mitochondria, and oxidative damage [11, 12].

Numerous studies have focused on the impact of acute/chronic ER stress on mitochondria bioenergetics, but less is known about the mediators of this ER-mitochondrial crosstalk [13, 14, 15]. In this study, we aimed to investigate the role of ROS accumulation in ER stress-induced changes in mitochondrial bioenergetics, biogenesis, and biodynamics, as described by mitochondrial respiration, mass/volume, and morphology. We examined those changes and the impact of ROS generation in human skeletal muscle myoblasts using the ER stress inducer tunicamycin and a synthetic antioxidant, Eukarion (EUK)-134, which has both superoxide dismutase and catalase activity [16].

## METHODS

### Cell culture and treatments

An immortalised human skeletal muscle cell line (donor age, 25 years) was provided as a gift to our group from the Institute of Myology, Paris [17]. Skeletal muscle cells were cultured in growth medium containing Dulbecco’s modified eagles medium (DMEM, Lonza, Nottingham, UK) and Medium-199 with Earle’s BSS (1:5, v/v) (Sigma-Aldrich, Dorset, UK), supplemented with 20% (v/v) heat inactivated foetal bovine serum (Gibco, Loughborough, UK), 1% (v/v) penicillin/streptomycin, 1% (v/v) L-glutamine (Lonza, Nottingham, UK), 10 μg/mL gentamicin, 25 ng/mL fetuin from foetal bovine serum, 0.2 μg/mL dexamethasone, 5 μg/mL recombinant human insulin (Sigma-Aldrich, Dorset, UK), 0.5 ng/mL recombinant human basic fibroblast growth factor, 5 ng/mL recombinant human epidermal growth factor, and 2.5 ng/ml recombinant human hepatocyte growth factor (Gibco, Loughborough, UK). Skeletal muscle myoblasts were incubated at 37°C in a humidified atmosphere of 5% CO_2_ until 80% confluence, and sub-cultured using 0.05% Trypsin/0.53 mM EDTA (1×) (Lonza, Nottingham, UK). Myoblasts were treated with the pharmaceutical ER stress inducer tunicamycin (0.1 μg/ml) in the absence or presence of EUK-134 (10 μM) for 24 hours. Data from our laboratory (not shown) and others have determined a dose of 10 μM EUK-134 to show efficacy in the absence of apoptosis/cell death - therefore, this concentration was chosen for this study [18].

### RNA isolation and Quantitative real-time polymerase chain reaction (PCR)

Following 24 hours of treatment, myoblasts were harvested using Dulbecco’s Phosphate-Buffered Saline (Lonza, Nottingham, UK) and stored at − 80°C. RNA of treated cells was isolated using EZ-RNA isolation kit (Biological Industries Beit Haemek, Israel) and cDNA was synthesised using iScript cDNA synthesis kit (Bio-Rad, UK). Real-time qPCR analyses were performed on a StepOnePlus™ Real-Time PCR System (Applied Biosystems) using QuantiNova SYBR Green PCR Kit (QIAGEN); 10 ng of cDNA per reaction was used in a total volume of 10 μl PCR reaction mixture. Three amplifications, consisted of an initial denaturation cycle at 95°C for 2 minutes, followed by 40 cycles of 20 seconds at 95°C (denaturation), 20 seconds at optimal annealing temperature, and 20 seconds at 72°C (extension), were performed for each primer, followed by a melt curve analysis. Threshold cycle for each target gene of interest was normalised to the housekeeping gene 18S (annealing temperature, 55.7°C), and analysed using the delta-delta (2^ΛΛCt^) method [19]. The sequences and annealing temperature of all mitochondrial- and ER stress pathway-associated primers used are provided in Supplementary Table 1 and 2, respectively.

### Measurement of mitochondrial bioenergetics using Seahorse Extracellular Flux Analyser

Real-time oxygen consumption rate (OCR) in skeletal muscle myoblasts was measured with a Seahorse XFp Extracellular Flux Analyser (Agilent Technologies, UK). Cells were seeded in an 8-well XFp cell culture microplate at a density of 7×10^3^ cells/well in 100 μl growth medium, and after adhesion, myoblasts were incubated with TN+/-EUK-134 under standard conditions. The Seahorse XFp Mito Stress Test was performed according to manufacturer’s instructions. Specifically, 1-hour prior the experiment, growth medium was replaced with unbuffered DMEM (pH 7.4) supplemented with 1 mM pyruvate, 2 mM L-glutamine, and 10 mM glucose, and the plate was equilibrated at 37°C in a non-CO_2_ incubator. Parameters of the cell bioenergetic phenotype were determined following the sequential addition of oligomycin (1 μM), carbonyl cyanide 4-(trifluoromethoxy) phenylhydrazone (FCCP, 2 μM), and rotenone/antimycin (0.5 μM). OCR and extracellular acidification rates, normalised using the bovine gamma globulin assay, were automatically calculated by Seahorse XFp software version 2.2.0.

### Quantification of mitochondrial morphology parameters using Confocal Microscopy

Skeletal muscle myoblasts were seeded on a 35-mm glass-bottom μ-Dish (ibidi^®^, Martinsried, Germany), treated as previously described, incubated with the cell-permeant MitoTracker Red CMXRos (5 μM, 30 minutes, 37°C) (Molecular Probes, Invitrogen, Paisley, UK) selective for living mitochondria, and fixed in 4% paraformaldehyde. Nuclei were counter-stained with 4’,6’-diamidino-2-phenylindole dihydrochloride (DAPI; 1/5000) (Sigma-Aldrich, Dorset, UK). Imaging was performed on a Leica TCS SP5 confocal microscope (Leica Microsystems, Milton Keynes, UK), using a 63×/1.4 oil immersion objective. Parameters of mitochondrial morphology were quantified using a macro on NIH ImageJ, created by Dagda et al. (2009) [20]. Briefly, following background subtraction and local contrast enhancement, region of interest (individual cell) was selected, and macro was activated to subject the images to threshold and transform them to binary.

### Measurement of ROS generation and oxidative damage markers

To quantify ROS generation and oxidative damage, skeletal muscle cells were seeded at 7×10^3^ cells/well in a black, clear bottom microplate (96 wells) and cultured in growth medium. After 24-hour treatment with TN+/-EUK-134, myoblasts were washed once with warm DPBS and incubated with different fluorophores in phenol red-free DMEM medium, in the dark at 37°C. Total intracellular ROS were determined using 2,7-dichlorofluorescein diacetate (DCFH-DA, 10 μM, 30 minutes) (Sigma-Aldrich, Dorset, UK). Intracellular and mitochondrial superoxide generation was measured using dihydroethidium (DHE, 5 μM, 20 minutes) (Sigma-Aldrich, Dorset, UK) and MitoSOX Red mitochondrial superoxide indicator (5 μM, 30 minutes) (Molecular Probes, Invitrogen, Paisley, UK), respectively. Following incubation with fluorophores, myoblasts were washed three times with DPBS and maintained in Seahorse assay medium. Endpoint fluorescence was measured using a Synergy™ multi-detection microplate reader (BioTek Instruments) with the following excitation and emission wavelength: DCFH-DA, 485/20 and 590/35 nm; DHE, 320/40 and 460/40 nm; and MitoSOX Red, 530/25 and 590/35 nm. Total protein thiols (sulphydryl) were quantified as per the manufacturer’s guidelines (Abcam, Cambridge, UK). Briefly, treated myoblasts were incubated with Thiol Blue sensor for 30 minutes on a shaker and samples were run through spin column. Absorbance was measured using a microplate reader at A280 nm and A680 nm.

For each experiment, the mean value derived from blank wells was subtracted to correct for background fluorescence/absorbance. All microplate reader measurements were normalised to total protein content per sample using the Pierce™ BCA Protein Assay and Bovine Gamma Globulin assay (Thermo scientific, Loughborough, UK).

### Assessment of mitochondrial membrane potential

Mitochondrial membrane potential ΔΨm) was interrogated using the JC-1 fluorophore (5,5’,6,6’-tetrachloro-1,1’,3,3’-tetraethylbenzimi-dazolylcarbocyanine iodide; 5 μM, 30 minutes, 37□C) (Abcam, Cambridge, UK), MitoTracker Red CMXRos (5 μM, 30 minutes, 37°C) (Molecular Probes, Invitrogen, Paisley, UK), or TMRM (Tetramethylrhodamine, methyl ester; 10 nM, 37°C) (Molecular Probes, Invitrogen, Paisley, UK). Following treatment, myoblasts were incubated with and maintained in TMRM solution, and endpoint fluorescence was read at excitation 530/25 nm and emission 590/35 nm. JC-1 normally forms red fluorescent aggregates. Red to green fluorescent shift occurs when mitochondrial membrane potential decreases, because of the presence of the green fluorescent monomeric form of JC-1 [21]. Following washing with DPBS, myoblasts were maintained in Seahorse assay medium and endpoint fluorescence from aggregate and monomer form was recorded on the microplate reader at excitation 530/25 nm and 485/20 nm, respectively, and emission 590/35 nm. Similarly, for MitoTracker Red CMXRos fluorescence staining, myoblasts were maintained in Seahorse assay medium and endpoint fluorescence was read at excitation 590/20 nm and emission 645/40 nm. Measurements were normalised to total protein content per sample using the Pierce™ BCA Protein Assay and Bovine Gamma Globulin assay.

### Measurement of mitochondrial mass/volume

Human skeletal muscle myoblasts were incubated with 100 nM MitoTracker Green FM (Molecular Probes, Invitrogen, Paisley, UK) prepared in a phenol red-free DMEM and incubated for 30 minutes at 37°C. Then, cells were washed once with DPBS and read by a fluorescence microplate reader (excitation 485/20 nm and emission 528/20 nm) or imaged using a LEICA DMI6000 B inverted microscope at 40× magnification (CTR6000 laser, Leica Microsystems). Fluorescence intensity was normalised to total protein content or nuclei number (using DAPI staining), respectively.

### Immunofluorescence staining

Human skeletal muscle myoblasts were washed twice with DPBS and fixed using 4% (w/v) paraformaldehyde. After 15 minutes fixation at room temperature, cells were washed and permeabilised using 0.5% (v/v) Triton X-100 for 15 minutes at room temperature. Then, cells were washed and blocked with 3% (v/v) goat serum supplemented with 0.05% (v/v) Tween-20 for 1 hour at room temperature. After washing, cells were incubated with rabbit anti-GRP78 (1/1000) (ab213258; Abcam, Cambridge, UK) at 4°C overnight. Secondary antibody conjugated with Alexa Fluor 488 (goat anti-rabbit IgG; 1/800; Invitrogen, Paisley, UK) and DAPI (1/5000; Sigma-Aldrich, Dorset, UK) were added to the cells for 1 hour at room temperature in the dark. Cells were maintained in DPBS and images were taken using a LEICA DMI6000 B inverted microscope at 20× magnification (CTR6000 laser, Leica Microsystems).

### Western blotting

Human skeletal muscle myoblasts were lysed using RIPA buffer protease/phosphatase inhibitors and the total protein in each sample was quantified using the Bovine Gamma Globulin assay. Twenty micrograms of protein were separated by 4-15% Mini-PROTEAN TGX Precast Gels (PAGE) (Bio-Rad, Hertfordshire, UK) and transferred to a nitrocellulose membrane. The membranes were blocked with 5% (w/v) fat free milk dissolved in Trisphosphate buffer with 0.0125% (v/v) Tween 20 for 1 hour and incubated with primary antibodies anti-GRP94 (ab3674), anti-MFN2 (ab56889), anti-SOD1 (ab13498), and anti-SOD2 (ab13533) (1/1000), as well as anti-β-actin (ab8226) (1/5000) as a loading control, at 4°C overnight. Secondary antibodies were added for 1 hour at room temperature and visualised using chemiluminescent substrate (Thermo scientific, Loughborough, UK) and LI-COR Odyssey Fc Imaging System (LI-COR Biosciences, Cambridge, UK).

### Statistical analyses

Data were assessed for normality of distribution by Shapiro-wilk test. Data assessed to be normally distributed were analysed using one-way ANOVA, with Tukey post hoc test. Data not normally distributed were analysed using Kruskal-wallis test, where appropriate. Data were analysed using GraphPad Prism version 8. A *p*-value ≤ 0.05 was considered to be statistically significant.

## RESULTS

### 1. ER Stress Pathway Activation

Significantly increased expression (fold change) of the UPR genes, *GRP78,* growth arrest and DNA damage-inducible gene 34 *(GADD34),* total X-box binding protein 1 *(XBP1),* cholesterol oxidase-peroxidase C/EBP homologous protein *(CHOP),* and ER-DnaJ-like 4 (*ERDJ4*), were observed in human skeletal muscle myoblasts treated with tunicamycin, as expected. Combination treatment with EUK-134 resulted in significantly attenuated expression of all genes, except from *GADD34* (**Figure 1A**). ER stress activation following tunicamycin treatment was further confirmed by significantly increased protein levels of *GRP94,* which, however, were not inhibited by EUK-134 (**Figure 1B**). GRP78 fluorescence intensity increased upon ER stress activation, however, there was no change in presence of EUK-134 (**Figure 1C and 1D**).

**Figure 1.**
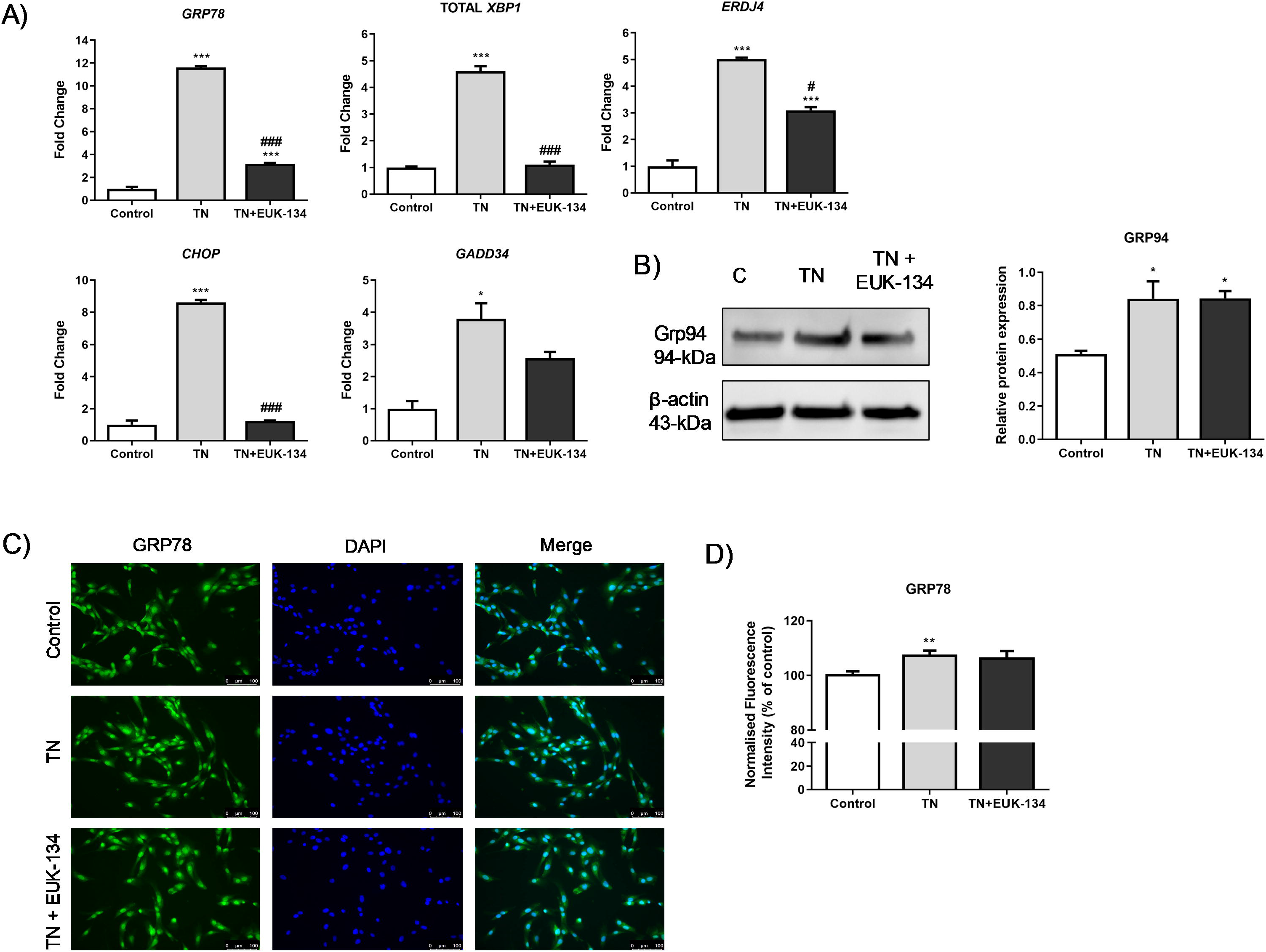
ER stress activation in tunicamycin-treated cells with or without EUK-134. **(A)** Relative fold changes in gene expression for ER stress markers. Data represent mean fold change normalised to 18S housekeeper gene ± SEM of ΔCt values (*n* = 3). **(B)** Representative western blot image and quantification of GRP94 protein levels relative to β-actin (loading control). Data represent mean ± SEM (*n* = 3). **(C)** Representative images of human skeletal muscle myoblast stained for GRP78 (green) and DAPI (blue). Images captured at 20× magnification. Scale bar = 100 μm. Brightness was adjusted equally in each image to enhance visualisation. **(D)** Quantification of total GRP78 fluorescence intensity normalised to nuclei number, relative to control (%). Data represent mean ± SEM, (*n* = 26) *\*p ≤* 0 .033, \*\**p* < 0.002, \*\*\**p* < 0.001 against vehicle control or tunicamycin alone (#).

### 2. Mitochondrial Oxygen Consumption and Mitochondrial Unfolded Protein Response

Tunicamycin-induced ER stress showed an overall increase in mitochondrial and non-mitochondrial respiration, which was attenuated in the presence of the antioxidant EUK-134 (**Figure 2A and 2D**). Basal OCR values were plotted versus the ECAR values to distinguish the metabolic profile of tunicamycin-treated myocytes in presence or absence of EUK-134, showing significantly increased basal OCR levels compared to the control group, with no change in basal ECAR levels (**Figure 2A and 2B**) [22]. EUK-134 was able to significantly inhibit tunicamycin-induced increase in basal respiration (**Figure 2B**). Decreased spare respiratory capacity (%, also called as reserve capacity) observed after ER stress induction was significantly increased following EUK-134 treatment, even compared to the vehicle control (**Figure 2E**). Consistently, depressed ATP-linked OCR was observed following tunicamycin treatment, (**Figure 2F**), which was further enhanced by a substantial increase in leak respiration, with more than half of oxygen consumed not being used for ATP production (**Figure 2G)**. Importantly, ER stress-induced proton leak was inhibited by EUK-134 treatment.

**Figure 2.**
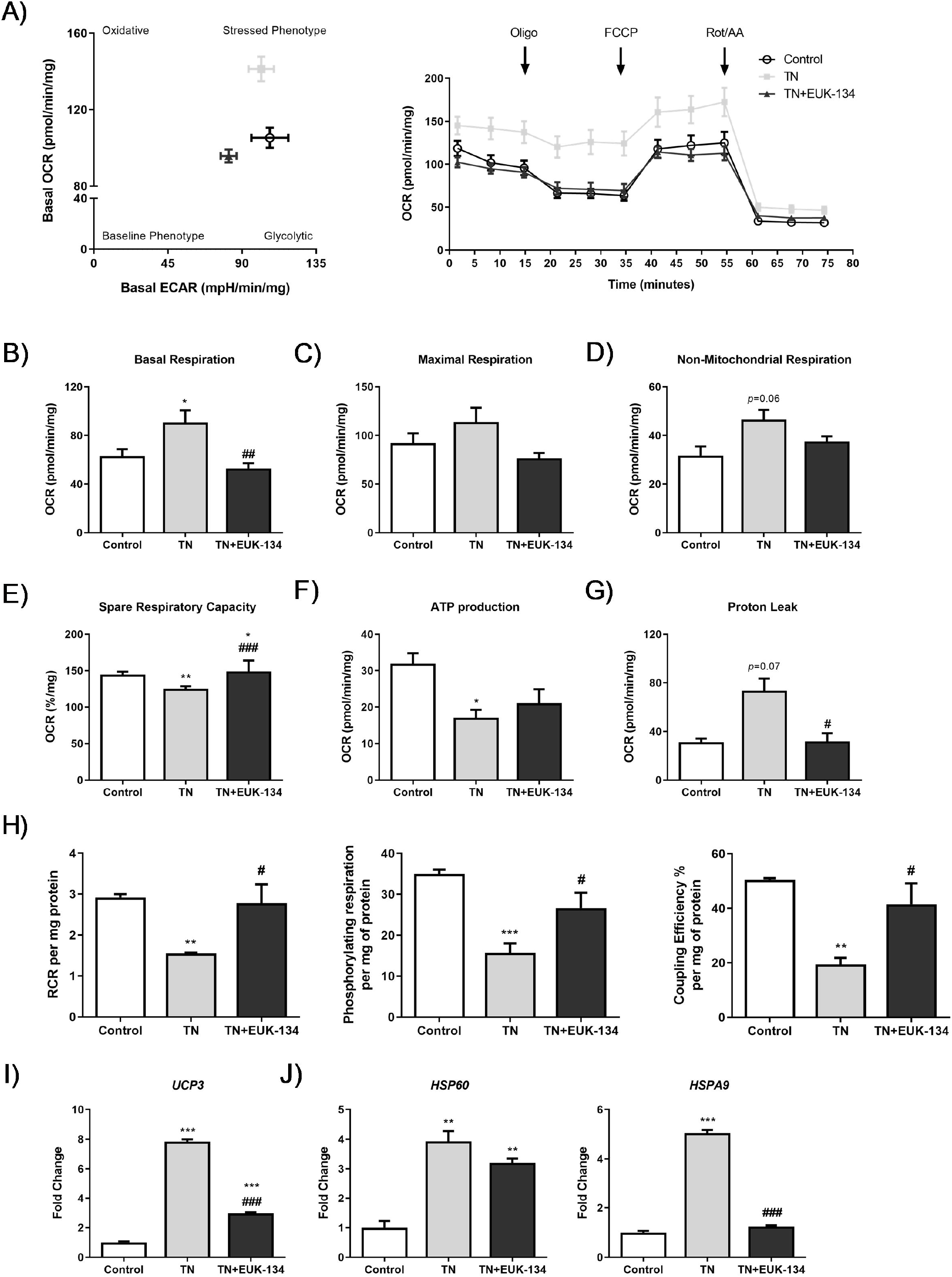
Cellular bioenergetics of tunicamycin-treated cells with or without EUK-134. **(A)** Bioenergetic status presented by basal oxygen consumption rate versus basal extracellular acidification rate, and real-time oxygen consumption rate measured under basal conditions followed by the sequential addition of oligomycin, FCCP, and rotenone/antimycin A mix. **(B–G)** Individual parameters of mitochondrial function; basal respiration, maximal respiration, non-mitochondrial respiration, spare respiratory capacity, ATP production, and proton leak, normalised to protein content. **(H)** Normalised respiratory flux control ratios, respiratory control ratio, phosphorylating respiration, and coupling efficiency (%), derived from the individual mitochondrial parameters. Data represent mean ± SEM, (*n* = 5) \**p* ≤ 0 .033, \*\**p* < 0.002, \*\*\**p* < 0.001 against vehicle control or tunicamycin alone (#). **(I)** Relative fold change in mRNA expression of *UCP3.* **(J)** Relative fold changes in mRNA expression of mitochondrial unfolded protein response markers, *HSP60* and *HSPA9.* Data represent mean fold change normalised to 18S housekeeper gene ± SEM of ΔCt values, (*n* = 3) \**p* ≤ 0 .033, *\*\*p* < 0.002, \*\*\**p* < 0.001 against vehicle control or tunicamycin alone (#). OCR, oxygen consumption rate; ECAR, extracellular acidification rate; RCR, respiratory control ratio.

To further evidence ER stress-induced mitochondrial dysfunction and investigate the impact of EUK-134, normalised respiratory flux control ratios were determined using the six parameters of mitochondrial function [21]. A significant decline was found in respiratory control ratio (RCR), phosphorylating respiration, and coupling efficiency under ER stress, which were inhibited by antioxidant treatment (**Figure 2H**). ER stress impaired the efficiency of mitochondrial respiration and decreased the potential ATP turnover leading to proton leak, and these effects were ROS-mediated. ROS-mediated decreased coupling efficiency and increased leak respiration, both indicative of proton leak-driven oxygen consumption, were also attributable to changes in the expression of the mitochondrial uncoupling protein 3 (*UCP3*), which were significantly decreased by EUK-134, but not to control levels (**Figure 2I**).

ER stress-induced activation of the mitochondrial unfolded protein response was also evident. Real-time qPCR results showed ROS-dependent increase in heat shock protein (HSP) Family A Member 9 (*HSPA9*) following tunicamycin treatment, but also ER stress-induced *HSP60* expression, which was upregulated in presence of EUK-134, as well (**Figure 2J**).

### 3. Mitochondrial Membrane Potential and Mitochondrial Mass

Changes in mitochondrial membrane potential following ER stress induction in presence or absence of EUK-134, as an additional marker of mitochondrial dysfunction, were next examined using different fluorophores. MitoTracker Red showed TN-induced hyperpolarisation, which was significantly inhibited by EUK-134 (**Figure 3A**). These results were also seen when assessing the accumulation of JC-1 polymers (red signal), indicative of increased number of hyperpolarised mitochondria (**Figure 3B**). Interestingly, JC-1 polymers to monomers ratio (red to green signal), showed TN-induced depolarisation which was prevented by EUK-134. This may be due to changes in mitochondrial mass or the existence of a pre-autophagic pool following asymmetrical mitochondrial fission [23, 24]. To examine whether the increase in MitoTracker Red and JC-1 polymers fluorescence intensity was attributable to a rise in mitochondrial mass/volume, TMRM was employed with or without MitoTracker Green to normalise to mitochondrial mass/volume. Indeed, TMRM showed mitochondrial hyperpolarisation after tunicamycin treatment (**Figure 3E**), while mitochondrial content normalisation produced a substantial loss mitochondrial membrane potential by tunicamycin with no changes observed in presence of EUK-134 (**Figure 3F**). In other words, tunicamycin induced increase in mitochondrial mass/volume that seems to have affected our initial interpretation; importantly this effect was inhibited by EUK-134 (**Figure 4A and B**). These results were further confirmed by increased expression (fold change) of *citrate synthase* and transcription factor A and mitochondrial precursor *(TFAM),* the initial enzyme of the citric acid cycle and a regulator of mitochondrial DNA transcription, respectively; both prevented by EUK-134. (**Figure 4C and D**). Collectively, data suggest that biogenesis is induced in response to ER and mitochondrial stress to compensate for changes in energetic demand.

**Figure 3.**
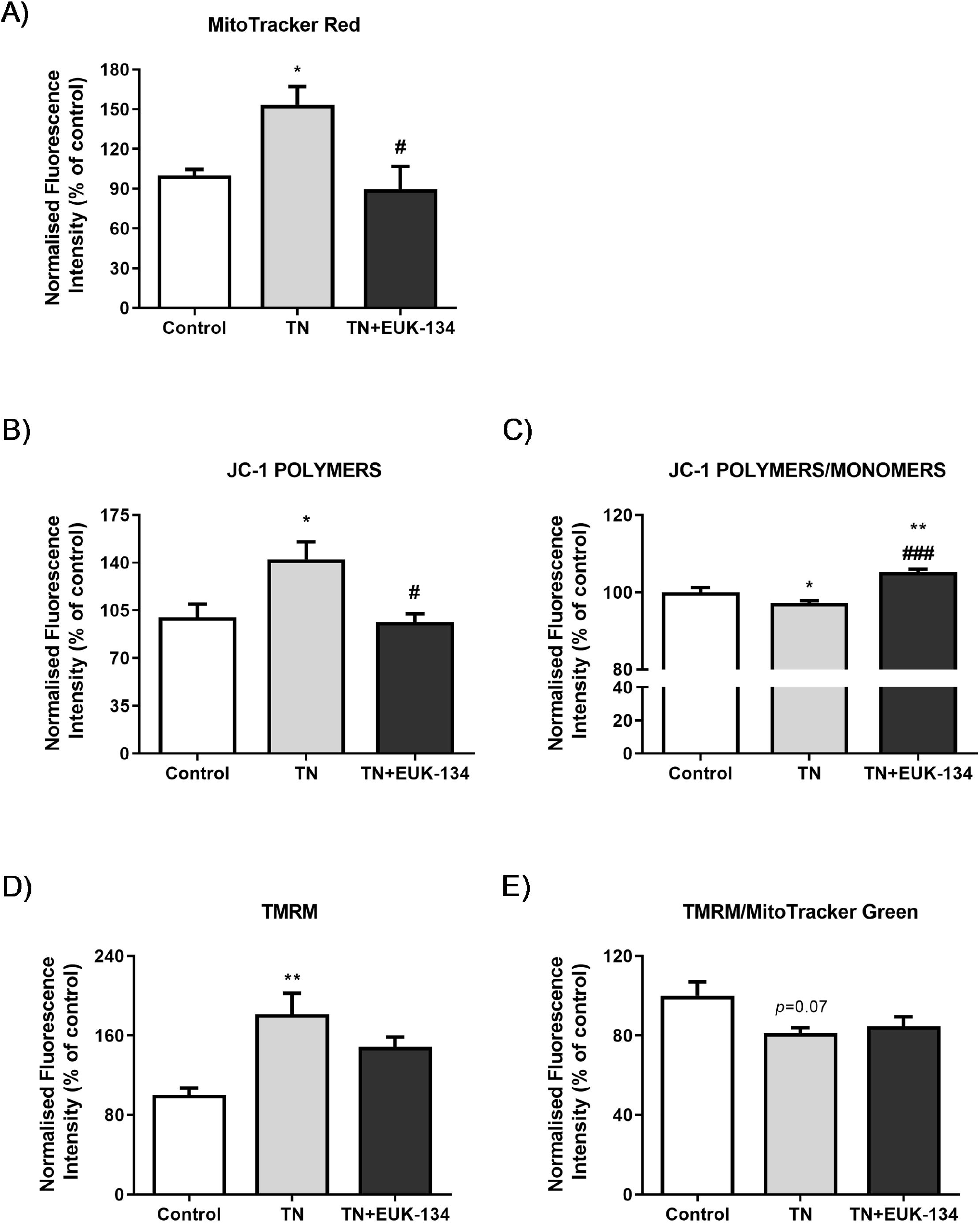
Mitochondrial membrane potential of tunicamycin-treated cells with or without EUK-134. Relative change (%) in **(A)** MitoTracker Red CMXRos fluorescence intensity (*n* = 4), **(B)** JC-1 red fluorescence (polymers) intensity (*n* = 8), **(C)** JC-1 red to green fluorescence (polymers/monomers) intensity (*n* = 8), **(D)** TMRM fluorescence intensity (*n* = 4), and **(E)** TMRM per MitoTracker Green (mitochondrial mass/volume) fluorescence intensity (*n* = 4), normalised to protein content. Data represent mean ± SEM, \**p* ≤ 0 .033, \*\**p* < 0.002, \*\*\**p* < 0.001 against vehicle control or tunicamycin alone (#).

**Figure 4.**
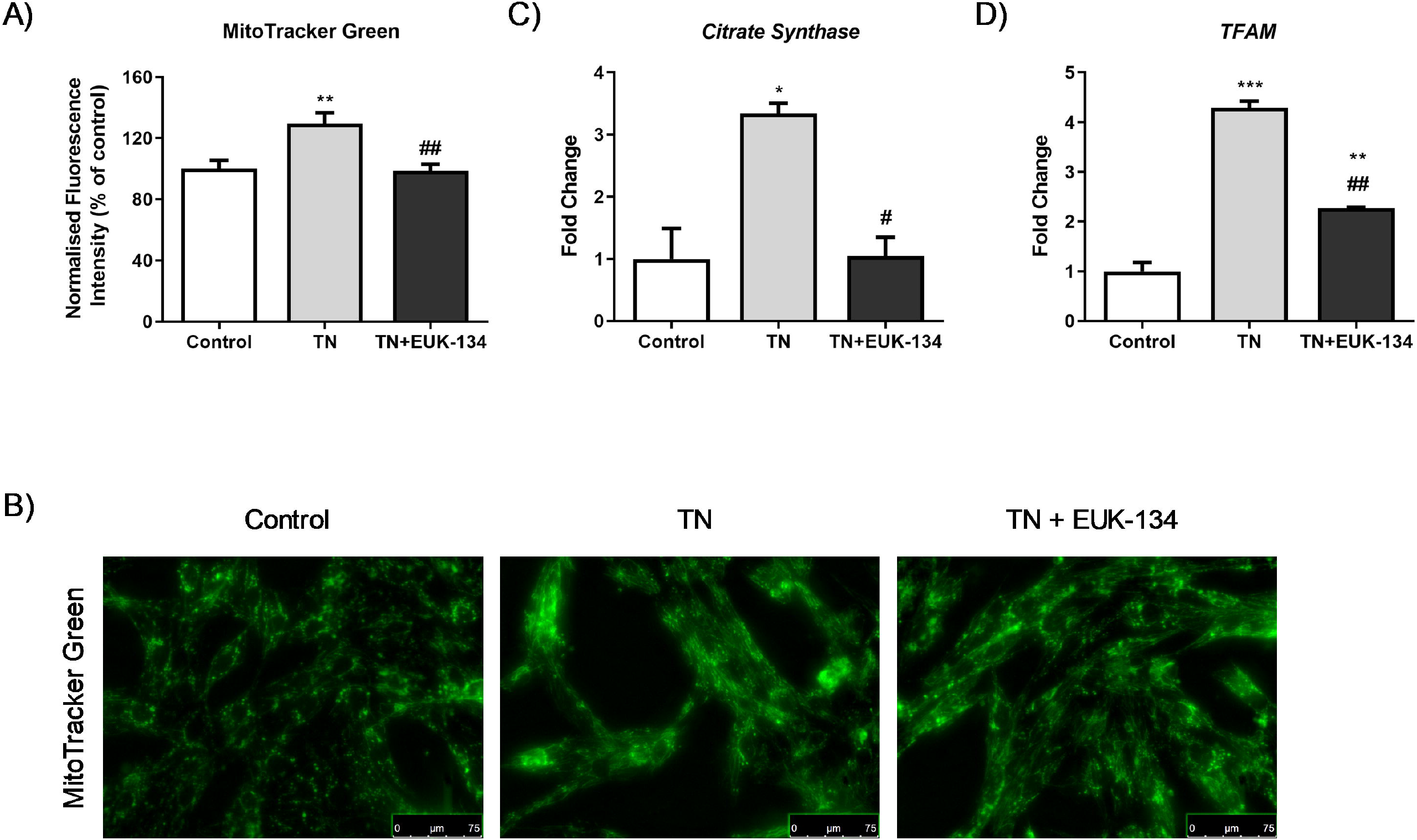
Mitochondrial biogenesis of tunicamycin-treated cells with or without EUK-134. **(A)** Representative graph of mitochondrial mass/volume showing the relative change (%) in MitoTracker Green fluorescence intensity, normalised to protein content. Data represent mean ± SEM, (*n* = 12) \**p* ≤ 0 .033, \*\**p* < 0.002, against vehicle control or tunicamycin alone (#). **(B)** Representative images of mitochondrial staining in human skeletal muscle myoblast using MitoTracker Green acquired under cell culture conditions. Images captured at 40× magnification. Scale bar = 75 μm. Brightness was adjusted equally in each image to enhance visualisation. **(C–D)** Relative fold changes in mRNA expression of *Citrate Synthase* and *TFAM.* Data represent mean fold change normalised to 18S housekeeper gene ± SEM of ΔCt values, (*n* = 3) \**p* ≤ 0 .033, \*\**p* < 0.002, \*\*\**p* < 0.001 against vehicle control or tunicamycin alone (#).

### 4. Mitochondrial Morphology: Fusion and Fission Events

Mitochondrial network structure was visualised using MitoTracker Red to explore mitochondrial dynamics, including fusion and fission processes, in response to ER stress and antioxidant intervention. Tunicamycin induced a significant increase in mitochondrial interconnectivity and elongation, indicative of mitochondrial fusion events, which can be associated with increased mitochondrial mass [25, 26]. These results were accompanying with an increase in the percentage of cytosol occupied by mitochondria. EUK-134 was able to drop TN-induced changes in mitochondrial interconnectivity and content to control levels, but not in elongation (**Figure 5A and B**). These findings were further confirmed by tunicamycin-induced upregulation of mitofusin-2 *(MFN2),* a gene associated with mitochondrial fusion; even though EUK-134 substantially decreased *MFN2* expression compared to tunicamycin treatment, it was still significantly upregulated compared to the control (**Figure 6A**).

**Figure 5.**
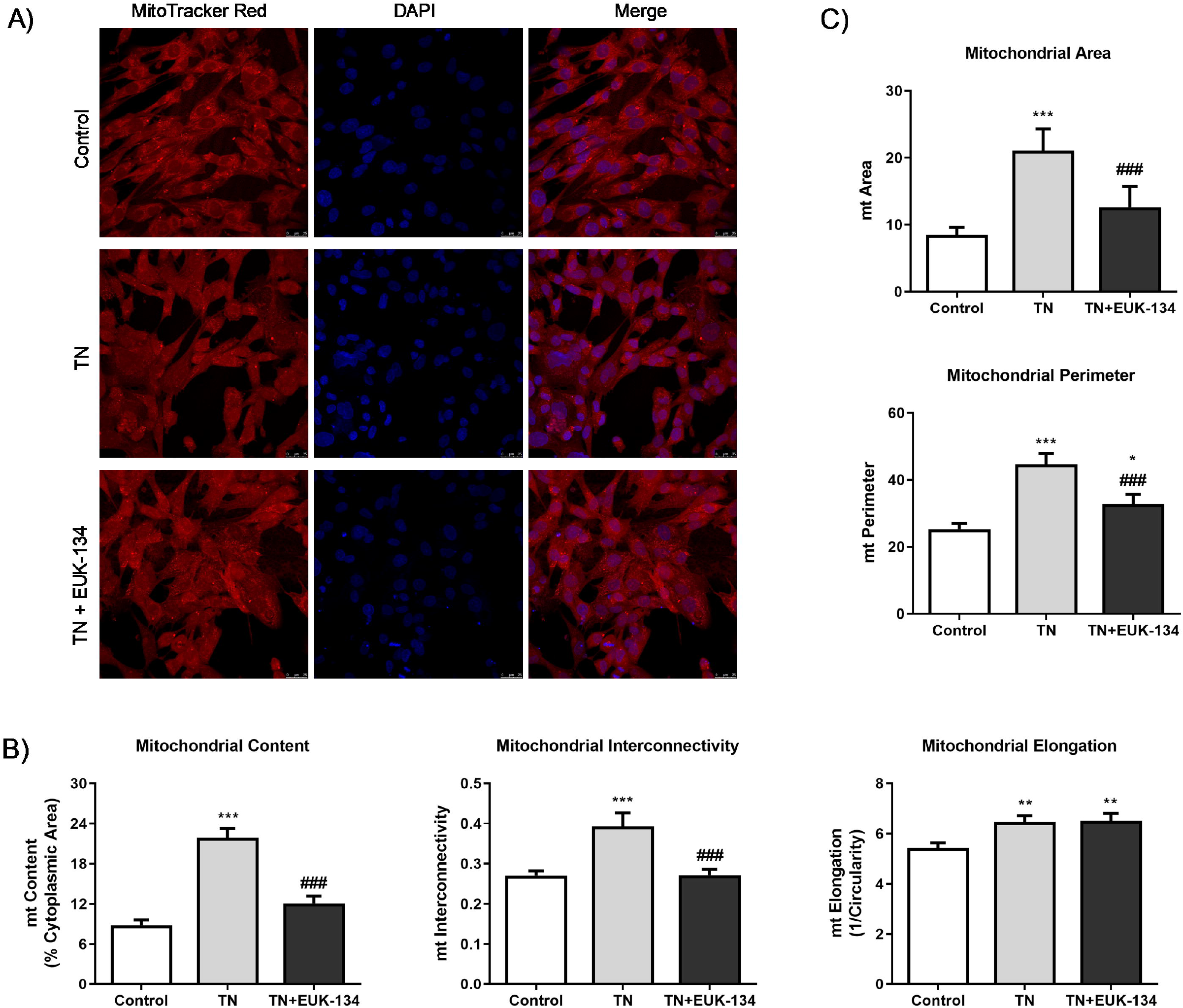
Mitochondrial morphological parameters of tunicamycin-treated cells with or without EUK-134. **(A)** Representative images of fixed human skeletal muscle myoblast stained with MitoTracker Red CMXRos. Images captured using 63×/1.4 oil immersion objective. Scale bar = 25 μm. **(B)** Mitochondrial content as % of cytoplasmic area, mitochondrial interconnectivity expressed by the ratio of mitochondrial area/perimeter, and mitochondrial elongation expressed by 1/circularity. **(C)** Mitochondrial fragmentation propensity described by mitochondrial area and perimeter. Data represent mean ± SEM, (*n* = 100 cells per group) \**p* ≤ 0 .033, \*\**p* < 0.002, \*\*\**p* < 0.001 against vehicle control or tunicamycin alone (#).

**Figure 6.**
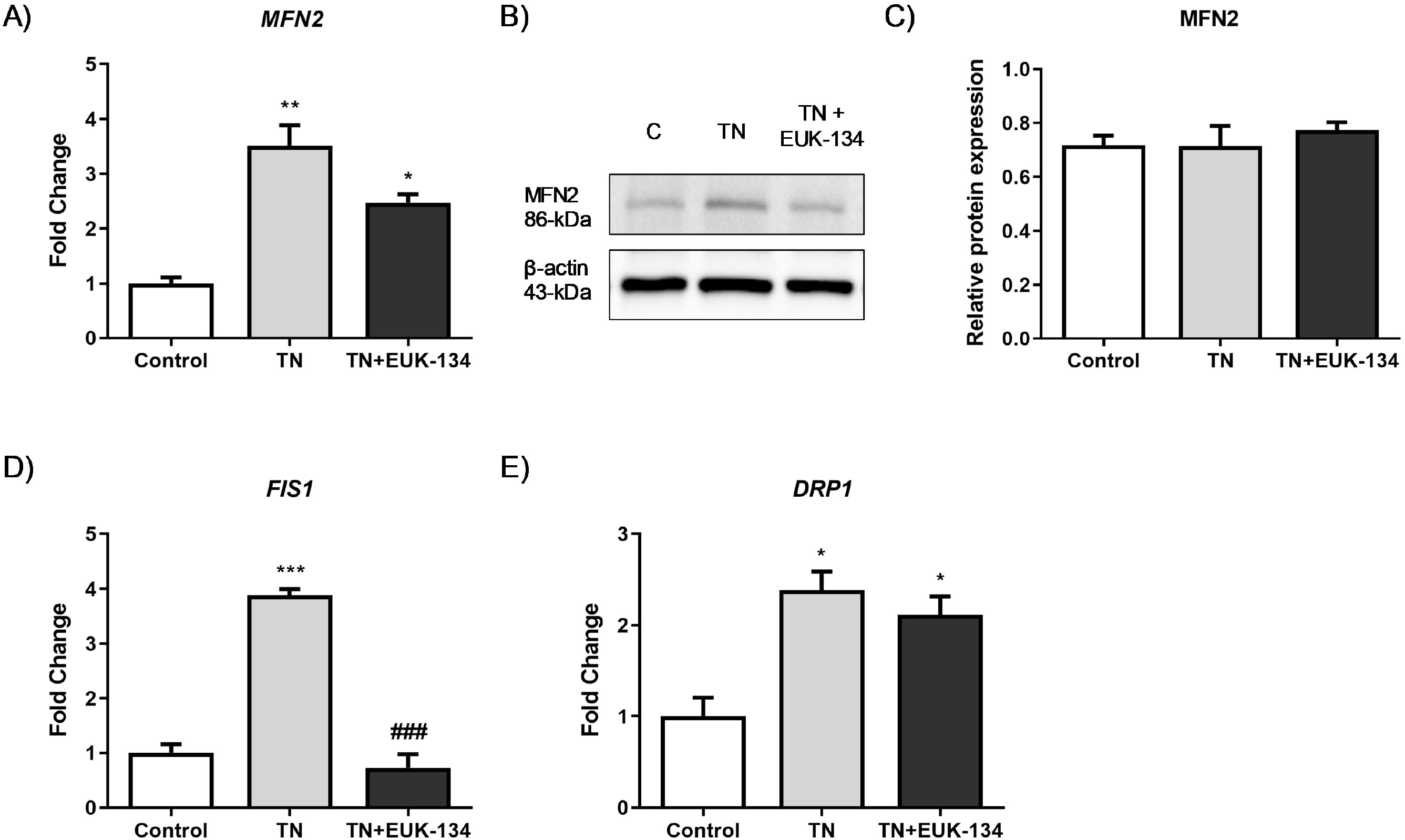
Mitochondrial dynamics of tunicamycin-treated cells with or without EUK-134. **(A)** Relative fold changes in mRNA expression of *MFN2* fusion-associated gene. Data represent mean fold change normalised to 18S housekeeper gene ± SEM of ΔCt values (*n* = 3) \**p* ≤ 0 .033, \*\**p* < 0.002 against vehicle control or tunicamycin alone (#). **(B–C)** Representative western blot image and quantification of MFN2 protein levels relative to β-actin (loading control). Data represent mean ± SEM (*n* = 3). **(D–E)** Relative fold changes in mRNA expression of *FIS1* and *DRP1* fission-associated gene. Data represent mean fold change normalised to 18S housekeeper gene ± SEM of ΔCt values (*n* = 3) \**p* ≤ 0 .033, \*\**p* < 0.002, \*\*\**p* < 0.001 against vehicle control or tunicamycin alone (#).

It has been described that mitochondrial perimeter and area are positively correlated with mitochondria about to undergo a fission event, which can result in increased mitochondrial number [25, 27]. MitoTracker Red staining showed increased tunicamycin-induced average mitochondrial perimeter and area, which were inhibited by EUK-134 (**Figure 5C**). These results are indicative of ROS-mediated fragmentation propensity in response to ER stress. This finding was further supported by increased expression changes in fission protein 1 *(FIS1)* and dynamin-related protein 1 *(DRP1)* genes, which are fission mediators. Overall, those results suggest that although the cell attempts to respond to ER stress-induced mitochondria dysfunction by increasing mitochondrial mass/volume, cells retain a propensity for fission.

### 5. ROS Generation

Since previous findings emphasised the importance of ROS in ER stressed-induced mitochondrial dysfunction, we aimed to further investigate ROS generation in our model. TN-treated cells exhibited higher levels of total cellular ROS (DCFH-DA fluorescence intensity) (**Figure 7A)** and specifically, superoxide levels (DHE fluorescence intensity), compared to the control group, which were decreased by EUK-134 treatment (**Figure 7B**). However, no changes were found on mitochondrial superoxide levels in presence of tunicamycin (**Figure 7C**), which might be because of its rapid conversion to hydrogen peroxide or peroxynitrite [11, 28, 29]. The observation of decreased thiols content is also indicative of elevated ROS generation and specifically, hydrogen peroxide release in the ER lumen through GSH oxidation to GSSG; EUK-134 was able to restore thiols levels (**Figure 7D**). We also examined the protein levels of superoxide dismutase (SOD) 1 and 2, as markers of ROS generation (**Figure 7E**). Results showed that tunicamycin induced a substantial increase in SOD1 as an adaptive response to ROS generation, but no changes observed in SOD2 protein levels (**Figure 7F and 7G**).

**Figure 7.**
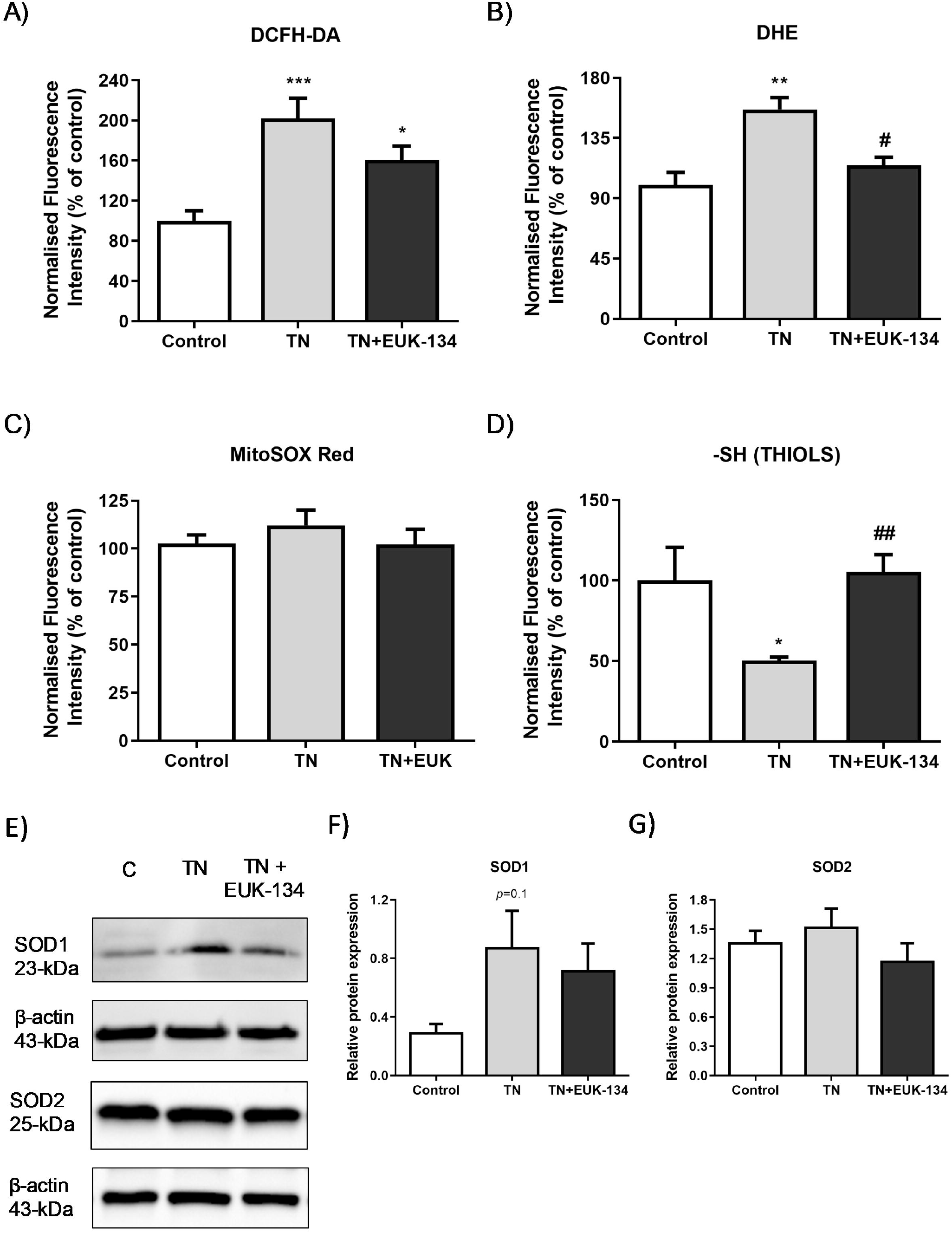
Markers of ROS generation in tunicamycin-treated cells with or without EUK-134. Relative change (%) in **(A)** total cellular ROS levels (DCFH-DA, *n* = 8), **(B)** total cellular superoxide levels (DHE, *n* = 4), **(C)** mitochondrial superoxide levels (MitoSOX Red, *n* = 20), and **(D)** total thiol content (*n* = 6), normalised to protein content. Data represent mean ± SEM, \**p* ≤ 0 .033, \*\**p* < 0.002, \*\*\**p* < 0.001 against vehicle control or tunicamycin alone (#). **(E–G)** Representative western blot image and quantification of SOD1 and SOD2 protein levels relative to β-actin (loading control). Data represent mean ± SEM (*n* = 4-5).

## DISCUSSION

Crosstalk between ER and mitochondria is a key cellular process and there is strong link between chronic ER stress and mitochondrial dysfunction [11]. Based on the existing evidence of ER stress activation as a mechanism involved in skeletal muscle weakness in patients with myositis [10], and the little knowledge on the role of ROS generation on ER stress downstream effects, this study aimed to determine the impact of the antioxidant EUK-134 in an *in vitro* model of ER stress-induced mitochondrial dysfunction in skeletal muscle, focusing on various aspects including mitochondrial function, biogenesis, and dynamics. EUK-134 was selected in the present study as it has been previously reported to exert beneficial effects on muscle atrophy and dysfunction induced by oxidative stress [30, 31, 32, 33].

Tunicamycin is an inhibitor of the *n-*glycosylation step of protein folding, leading to accumulation of unfolded glycans within the ER [34]. Tunicamycin has been extensively used in numerous studies to activate ER stress pathway in mouse and human skeletal muscle cells [35, 36, 37]. In this study, tunicamycin-induced ER stress, validated by an increase in UPR pathway markers, was partially ameliorated by EUK-134, with inhibitory effects on mRNA levels of *GRP78, ERDJ4,* and *total XBP1.* Even though the dose of tunicamycin was relatively lower compared to other studies [13, 37, 38], the 24-hour incubation has been used to exceed the early phase of ER stress [13, 39], and the expression of *CHOP*, a pro-apoptotic transcription factor, as a marker of prolonged/maladaptive ER stress [40]. Importantly, this study showed that EUK-134 diminished ER stress-induced upregulation of *CHOP* mRNA expression, protecting the cells from apoptosis initiation.

Previous studies have reported an increase in mitochondrial respiration and mass as an adaptive response to ER stress, which when unresolved, leads to mitochondrial dysfunction and cell death [13, 26, 39]. A major finding of this study is that prolonged tunicamycin administration induced impaired mitochondrial function that was mediated by ROS generation. In agreement with previous findings, prolonged tunicamycin treatment promoted basal mitochondrial respiration upon ER stress [14], which can be considered as an attempt of the cell to meet ATP demand [41], as they showed inability to shift to glycolysis. However, elevations in OCR were not correlated with ATP production, as previously described, with total cellular ATP levels substantially reduced under prolonged ER stress (20 hours, 0.5 μg/ml) compared to early ER stress (1-4 hours) and the control group [13]. Diminished ATP turnover and impaired oxidative phosphorylation were also evident by suppression of reserve capacity, coupling efficiency, RCR, and phosphorylating respiration induced by prolonged ER stress, showing the inability of the cell to respond to energetic demands. Importantly, those changes induced by ER stress were mitigated by EUK-134, highlighting the important role of ROS generation in ER stress-induced mitochondrial dysfunction. Increased basal respiration can also be associated with other sources of oxygen consumption, including ROS generation [41]. In accordance with this, the present study showed substantial rise in non-mitochondrial OCR upon ER stress, which was further supported by elevated changes in the markers of oxidative stress. Specifically, tunicamycin increased total cellular ROS generation, including superoxide levels, and reduced thiol content, which were inhibited by EUK-134. However, no changes were noticeable in mitochondrial superoxide levels. This finding can be likely explained by the presence of nitric oxide production that has been previously shown to compete with superoxide dismutase and reduce superoxide availability to produce peroxynitrite [28, 42]. Further supporting this hypothesis, a study on prostate cancer cells revealed that prolonged ER stress induced by tunicamycin, is highly correlated with endothelial nitric oxide synthase upregulation and nitric oxide production [43]. A previous study has shown that EUK-134 can reduce nitric oxide, and subsequently, peroxynitrite production in proximal tubular cell injury [44]. It is also known that thiols are oxidised by peroxynitrite into disulphide bonds [45]. Consistently with these findings, the present study showed that EUK-134 increased thiol content under ER stress conditions, potentially by inhibiting peroxynitrite production.

Mitochondrial dysfunction with noticeable increase in proton leak, as observed in the current study, has been correlated with changes in mitochondrial membrane potential [46]. Initial investigation showed tunicamycin-induced mitochondrial membrane hyperpolarisation. However, a previous study has emphasised the importance of normalising this parameter to not only protein content, but also, mitochondrial content [24]. Normalisation to mitochondrial mass/volume showed mitochondrial membrane depolarisation under ER stress, supporting its effects on mitochondrial biogenesis. Specifically, this study showed stimulation of mitochondrial biogenesis by tunicamycin as a response to increased ATP demands induced by ER stress, which was not noticeable in presence of EUK-134. This finding was consistent with another study showing increased mitochondrial mass in an animal model of mitochondrial myopathy, induced by respiratory chain deficiency [47].

Previous studies have shown a distinct impact of oxidative stress on mitochondrial structural network. In consistent with our study, high respiration rates but decreased reserve capacity, as well as loss of mitochondrial membrane potential were induced by hydrogen peroxide and preceded mitochondrial fragmentation in mouse skeletal muscle myocytes [48]. Specifically, it has been found that prior to mitochondrial fragmentation, mitochondria elongate and fuse as an adaptive response to cellular insults, including ER stress and ROS generation [26, 49]. Changes in mitochondrial structure have shown to determine cell fate when autophagy is stimulated, with elongated mitochondria being able to escape from degradation and maintain cell viability [50]. Similarly, in our model, it was found that 24 hours incubation with tunicamycin maintained fusion processes, which were partially suppressed by EUK-134. Fusion processes were described by elongated and interconnected mitochondria, as well as upregulated *MFN2* mRNA expression, which can be positively correlated with increased mitochondrial mass [25]. Taken together, these findings suggest ROS-mediated increase in mitochondrial fusion events and accumulation of dysfunctional mitochondrial mass under maladaptive ER stress conditions.

Since it is known that mitochondrial dysfunction precedes mitochondrial fragmentation, we examined the propensity of mitochondria to fragment in our ER stress model, as suggested by Westrate et al. [27]. Tunicamycin increased morphological parameters predictive of future mitochondrial fragmentation, including perimeter, the most prominent one, as well as mitochondrial area. These results were further supported by our findings of increased changes in mRNA levels of fission markers, *FIS1* and *DRP1.* Importantly, those events were mitigated by EUK-134, indicating that ER stress-associated fragmentation events are mediated by ROS. ROS-induced mitochondrial depolarisation and fragmentation also showed stimulation of ER UPR that further activated mitochondrial UPR in mouse skeletal muscle myoblasts [51]. These findings are in agreement with our study that showed TN-induced expression of *HSP60* and *HSPA9* mitochondrial UPR mediators, with *HSPA9* being completely inhibited by EUK-134.

## CONCLUSION

EUK-134 can protect against aspects of ER stress-induced mitochondrial dysfunction, biodynamics, and biogenesis in human skeletal muscle cells, highlighting a role of ROS in instances of prolonged ER stress, such us in myositis. Overall, this work provides a possibility of quenching ROS generation as an avenue for beneficial impact for ER stress-related diseases.

## ACKNOWLEDGMENTS

This study was funded by a Faculty PhD Studentship awarded to AT by The Manchester Metropolitan University.

