## Supplemental Table 1 for "Eukarion-134 attenuates endoplasmic reticulum stress-induced mitochondrial dysfunction in human skeletal muscle cells"

| **Table 1. The sequences and annealing temperature of mitochondrial-associated primers** | | | |
| --- | --- | --- | --- |
| **Target mRNA** | **Annealing Temperature (°C)** | **Forward Primer Sequence**  **(5'-3')** | **Reverse Primer Sequence**  **(5'-3')** |
| *MFN2* | 58 | AGTTGGAGCGGAGACTTAGC | ATCGCCTTCTTAGCCAGCAC |
| *HSP60* | 58 | GAACAGCTAACTCCAAGTCAGA | CAGCCGCTCTGAGAACTTCA |
| *TFAM* | 58 | CTGCACTCTGTCCCTCACTC | GGGTAACCGAAGCATTTCTGC |
| *DRP1* | 58 | TCACCCGGAGACCTCTCATT | TCTGCTTCCACCCCATTTTCT |
| *Citrate Synthase* | 58 | TGATGAGGGCATCCGTTTCC | GTTCTTCCCCACCCTTAGCC |
| *FIS1* | 58 | AGGCCTTAAAGTACGTCCGC | TGCCCACGAGTCCATCTTTC |
| *UCP3* | 55.7 | GGGTCAACCTGGGATGTAGC | TCCCTAACCCTCCCCATCAG |
| *HSPA9* | 58 | AGAAGACCGGCGAAAGAAGG | TGTTGCACTCATCAGCAGGT |
| **Abbreviations:** MFN1, mitofusin 2; HSP60, heat shock protein 60; TFAM, transcription factor A, mitochondrial precursor; DRP1, dynamin related protein 1; FIS1, fission 1; UCP-3, uncoupling protein 3; and HSPA9, heat shock protein family A member 9. | | | |
