## Supplemental Table 2 for "Eukarion-134 attenuates endoplasmic reticulum stress-induced mitochondrial dysfunction in human skeletal muscle cells"

| **Table 2. The sequences and annealing temperature of endoplasmic reticulum stress-associated primers** | | | |
| --- | --- | --- | --- |
| **Target mRNA** | **Annealing Temperature (°C)** | **Forward Primer Sequence**  **(5'-3')** | **Reverse Primer Sequence**  **(5'-3')** |
| *GRP78* | 59.3 | TGACATTGAAGACTTCAAAGCT | CTGCTGTATCCTCTTCACCAGT |
| *Total XBP1* | 59.3 | GGCATCCTGGCTTGCCTCCA | GCCCCCTCAGCAGGTGTTCC |
| *ERDJ4* | 59.3 | TCGGCATCAGAGCGCCAAATCA | ACCACTAGTAAAAGCACTGTGTCCAAG |
| *CHOP* | 59.3 | GGAGCATCAGTCCCCCACTT | TGTGGGATTGAGGGTCACATC |
| *GADD34* | 59.3 | CCCAGAAACCCCTACTCATGATC | GCCCAGACAGCCAGGAAAT |
| **Abbreviations:** GRP78, glucose-regulated protein 78 kDa; total XBP1, total X-box binding protein 1; ERDJ4, ER-DnaJ-like 4; CHOP, cholesterol oxidase-peroxidase C/EBP homologous protein; and GADD34, growth arrest and DNA damage-inducible gene 34. | | | |
